# A Reassessment of the Taxonomic Position of Mesosaurs Based on Two Data Matrices

**DOI:** 10.1101/391813

**Authors:** Michel Laurin, Graciela Piñeiro

## Abstract

The Early Permian mesosaurs are the oldest known primarily aquatic amniotes. Despite the interest that they have generated over time, their affinities remain controversial. Recently, two hypotheses have been supported, in which mesosaurs are either the sister-group of all other sauropsids, or the sister-group of other parareptiles. We recently upheld the former hypothesis, but in the latest study on mesosaur affinities, MacDougall et al. published a study highly critical of our work, while upholding the hypothesis that mesosaurs are basal parareptiles. We expect that the debate about mesosaur affinities will continue in the foreseeable future, but we wish to respond to the two central comments published by MacDougall et al. in 2018, who argue that variability in the temporal fenestration of early sauropsids, combined with the omission of several recently-described parareptile taxa, explain the differences in topologies between their study and ours. Reanalyzing our data matrix and theirs without characters linked with temporal fenestration, and removing from their matrix the parareptile taxa that they added (and that we omitted) does not alter the resulting topologies. Thus, their main conclusions are false; the differences in taxonomic position of mesosaurs must result from character choice and scoring differences.

## 3 Introduction

Mesosaurs, known from the Early Permian of southern Africa, Brazil and Uruguay, are the oldest known amniotes with a primarily, though probably not strictly, aquatic lifestyle (Nuñez Demarco et al., 2018). Despite having attracted attention of several prominent scientists, such as Wegener (1966), who used them to support his theory of continental drift, and the great anatomist and paleontologist von Huene (1941), who first suggested the presence of a lower temporal fenestra in *Mesosaurus*, several controversies still surround mesosaurs. One concerns the presence of the lower temporal fenestra in mesosaurs, which we support (Piñeiro et al., 2012a; Laurin and Piñeiro, 2017:4), contrary to Modesto (1999; 2006) and MacDougall et al. (2018); the other concerns the systematic position of mesosaurs, which have been argued, in the last decades, to be either the basalmost sauropsids (Laurin and Reisz, 1995; Laurin and Piñeiro, 2017), or the basalmost parareptiles (Gauthier et al., 1988; Modesto, 1999; MacDougall et al., 2018), if we use a branch-based definition of Parareptilia as Laurin and Reisz (1995) did, but using *Procolophon trigoniceps* as an internal specifier rather than turtles, because of the controversy surrounding the affinities about turtle origins.

In their recent response to our recent paper on the taxonomic position of mesosaurs, MacDougall et al. (2018) make a number of problematic claims, which we wish to discuss. These claims are that we used an outdated matrix and ignored over two decades of parareptile research, that our taxon selection was insufficient and that along with variability in temporal fenestration in parareptiles, all these choices explain the different taxonomic position of mesosaurs that we obtained (as the basalmost sauropsids rather than the basalmost parareptiles). Below, we respond to these claims by providing additional background data and by performing various analyses of their matrix and ours that show, among other things through taxon and character deletion, that neither the omission of some taxa, nor variability in temporal fenestration explains the differences in topologies between our study and theirs. We also highlight problems with their analyses and discuss why reusing phenotypic data matrices elaborated by other systematists is difficult.

## 3 Materials and Methods

The reanalyses below use our data matrix (Laurin and Piñeiro, 2017), on which we deleted characters linked to temporal fenestration, to assess the impact of this complex of characters on the resulting topology. We also reanalyze the data matrix of MacDougall et al. (2018), both unmodified, and modified by ordering characters that form morphoclines, and modified further by removing the parareptile taxa found in the matrix of MacDougall et al. (2018) but not in ours (Laurin and Piñeiro, 2017). We have not studied the scores of the cells of the data matrix of MacDougall et al. (2018) because this would be very time-consuming and would largely duplicate a more ambitious project focusing on early amniote phylogeny and the origin of turtles (mesosaurs are in the matrix, though the project does not focus particularly on their affinities) undertaken by one of us (ML) in January 2018 in collaboration with Ingmar Werneburg, Gabriel Ferreira and Marton Rabi.

All phylogenetic analyses were carried out with PAUP (Swofford, 2003) version 4.0a, build 163 for Macintosh (the latest version available as of July 28, 2018) using the branch and bound algorithm for our own data matrix, and using the heuristic search with 50 random addition sequence replicates, holding 3 trees at each step, and using the tree-bisection-reconnection (TBR) algorithm for the various versions of the matrix by MacDougall et al. (2018) because that matrix had too many taxa to use the branch and bound algorithm. The branch and bound algorithm guarantees to find all the most parsimonious trees. No heuristic algorithm provides similar guarantees, but we verified every time that all tree islands of most parsimonious trees had been recovered at least three times (most were recovered far more frequently and all were found at least 4 times) to be reasonably certain that we had all the most parsimonious trees. These allow us to test the main claims made by MacDougall et al. (2018).

We make all the new data (modified list of characters, in which we document which characters were ordered, and how the states had to be reordered to reflect the underlying continuous characters, and modified Mesquite Nexus file, which incorporates these ordering modifications) available on the journal web site.

## 4 Results

### 4.1 Taxon selection in recent studies on mesosaur affinities

MacDougall et al. (2018) claim that we « use of an outdated phylogenetic matrix and the fact that the authors patently ignore over two decades of parareptilian research ». This claim about two closely related points is factually wrong. The second point (that we ignore two decades of parareptilian research) is refuted by a simple examination of the bibliography of our paper. We did not claim to have cited all recent papers on parareptiles (given that our paper was not a review of recent studies on parareptiles), but we cited several papers published in the 1998–2018 period that have a high parareptiles content (Reisz and Scott, 2002; Cisneros et al., 2004; Müller and Tsuji, 2007; Modesto et al., 2009; Lyson et al., 2010, 2013; Tsuji et al., 2010, 2012; Lee, 2013; Bever et al., 2015), not including those about mesosaurs, given that our results suggest that mesosaurs are not parareptiles. The first point (that the matrix is obsolete) is equally factually wrong as shown by the fact that we updated extensively the original matrix (Laurin and Reisz, 1995) using both the literature and direct observations of specimens, especially of mesosaurs, and we explained this clearly (Laurin and Piñeiro, 2018:4). For instance, *Owenetta,* which we added to the matrix (it was not in the matrix of Laurin & Reisz, 1995) was scored using the detailed description of Reisz and Scott (2002). We also added one of the basalmost, best-known parareptiles (*Acleistorhinus*), to test better previous hypotheses that mesosaurs are basal parareptiles, and we also made other changes to the taxon set (see below). Thus, given all these changes, we do not consider that we used an outdated phylogenetic matrix.

MacDougall et al. (2018) object the use of suprageneric taxa as terminals because they claim that the resulting higher rate of polymorphism can weaken support values. While the increased polymorphism in OTUs corresponding to large clades is unavoidable and simply reflects reality, the relationship between that rate of polymorphism and support values in a matrix are far more complex than suggested by MacDougall et al. (2018). This is illustrated by the fact that the taxa (typically ranked as genera) that they introduced to replace the OTU Synapsida yields a tree that suggests that this taxon, as currently delimited, is paraphyletic! Of course, given the phylogenetic definition of Synapsida that has been proposed under the PhyloCode, Synapsida would remain monophyletic but varanopids and the presumed ophiacodontid *Archaeothyris*, in their most parsimonious trees (and under both of their analyses), move out of Synapsida and become basal sauropsids. Thus, the support value of Synapsida on their tree (not provided) is presumably very low (bootstrap values) to negative (Bremer or decay index), depending on which index is used to assess it. This does not negate the value of the addition of these small OTUs to replace Synapsida because the new hypothesis about the position of varanopids (outside Synapsida) is an interesting result. Thus, as epitomized in “A hitchiker's guide to the galaxy” (Adams, 2017), it is more important to have a good question than a precise answer, and in phylogenetics, it is perhaps more important to get interesting results that raise new questions (about the monophyly of Synapsida and Diapsida as currently delimited, for instance) than to obtain high support values, which, in some cases, may simply reflect the fact that only a trivial question was asked (about affinities between a small set of closely related taxa, for instance). We also moved (partly) in that direction (selecting smaller terminal taxa) by breaking up the Testudines OTU present in Laurin and Reisz (1995) into *Odontochelys, Proganochelys* and Chelonii, which could be broken up further in subsequent studies. Note also that there is no special justification for using nominal genera as OTUs given the subjective nature of Linnaean categories; all taxonomic ranks, even species, are artificial constructs, ontologically empty designations (Ereshefsky, 2002). In addition, all taxa represent phylogenetic hypotheses, so the most rigorous approach would be to do a specimen-level analysis, though this would result in much more missing data than working on taxa, and this raises new problems (e.g., Simmons 2012 a, b).

### 4.2 Phylogeny and evolution of Permo-Carboniferous amniotes

The findings by MacDougall et al. (2018) that Synapsida, as defined under the draft PhyloCode (Laurin and Reisz, in press) may exclude varanopids and the ophiacodontid *Archaeothyris* (other ophiacodontids were not included), which appear at the base of Sauropsida in their trees, raises two important points that MacDougall et al. (2018) did not discuss. First, it now seems likely that the phylogeny of Permo-Carboniferous amniotes is much less robust than previously thought, given that our analysis also found unorthodox results (parareptiles nested within diapsids). This is also highlighted by the fact that protorothyridids (represented by *Paleothyris* and *Protorothyris*), on their consensus trees, are paraphyletic.

Second, their topology, if correct, implies that the lower temporal fenestra is an amniote synapomorphy, a possibility that we suggested earlier (Piñeiro et al., 2012a) on the basis of the presence of a fenestra in mesosaurs. MacDougall et al. (2018) clearly viewed this part of their tree as preliminary and problematic, but it is a result that should be investigated further, as they pointed out, and it adds some support to the hypothesis that lower temporal fenestra is an amniote synapomorphy. They also discussed fenestration extensively and suggested that its great variability in amniotes decreased it taxonomic value, a point that we already made (Piñeiro et al., 2012a), and suggested that it raised doubts about our results. This last point is misleading because our matrix includes only two characters directly linked to temporal fenestration (characters 30 and 31). MacDougall et al. (2018), in their first analysis, emphasized temporal fenestration much more than we did by adding four characters linked to temporal fenestration (their characters 171–174, even though the older version of their matrix already had three other characters also linked with temporal fenestration (their characters 44–46). They removed these characters (along with our two temporal fenestration characters, which they also had) in their second analysis to assess the impact of fenestration on their results; this impact is apparently negligible, which does not support their claim.

MacDougall et al. (2018) criticized us for accepting our own anatomical interpretations (rather than theirs) about the temporal fenestra of mesosaurs: “the authors adhere to the interpretation of Piñeiro et al. (2012[a]) that *Mesosaurus* possessed a lower lateral temporal fenestra, a condition that actually may be absent or ontogenetically variable within the taxon”. This is a strange comment given that science is based on observation rather than authority, and that we have had access to far more specimens than them to support our interpretations. Should scientists prefer other's opinions over their own observations? This would run counter to the most basic scientific principles. We have seen no good evidence of the absence of the fenestra in any mesosaurid so far.

They also claim that “specimens with supposed temporal fenestration, such as that presented in Piñeiro et al. (2012[a]), are extremely poorly preserved”. This is wrong and misleading. Our specimens preserve bone (Fig. 1 C–E), whether the specimens studied by Modesto lacked bone (Modesto, 2010: 347): “All specimens of *Mesosaurus tenuiden*s examined here are preserved as natural moulds in black shale. These were cast in latex rubber and drawn from photographs or by use of a camera lucida.” Thus, we believe that at least some of our specimens are better-preserved than those described by Modesto, which is not surprising given that the Mangrullo formation, from which most of our specimens originate, is a recognized Konservat-Lagerstätte (Piñeiro et al., 2012b). No author of MacDougall et al. (2018) saw more than a small part of the specimens that we have studied, so they are not in a good position to discuss preservation of the specimens that support our interpretation. In any case, quality of preservation can be assessed through several criteria, such as whether or not bone is present, the degree of flattening of the skeleton, and whether or not elements are broken and in articulation. According to three of these four criteria (presence of bone, articulation and whether or not bones are broken), our specimens are better than those studied by Modesto (2006) because in the latter, the temporal region is disarticulated, bone is absent (only impression remains), and several elements appear to be incompletely preserved, with broken edges, as shown by the high variability of the shape of the squamosal in the specimens illustrated by Modesto (2006). Also, disarticulation of the specimens studied by Modesto is such that the basisphenoid is visible in dorsal view (Modesto, 2006: fig. 3), or the epipterygoid is visible in lateral view (Modesto, 2006: fig. 6), and in the latter specimen, it is obvious that the jugal is bifurcate posteriorly and that it defined the anteroventral corner of the lower temporal fenestra, though Modesto (2006: 352) interpreted this region differently. Modesto (2006) argued that the squamosal has a complementary shape and overlapped the posterior edge of the jugal, but given the extreme variability in the preserved portion of the squamosals illustrated by Modesto (2006), this interpretation seems to rest on tenuous evidence. For the other preservation criterion (flattening), quality is equivalent between the specimens from Uruguay and those from Brazil and South Africa studied by Modesto (2006, 2010). One last point to consider is that the Mangrullo formation of Uruguay has yielded isolated elements, including those that border the temporal fenestra (Fig. 1A, B), and these support our interpretation.

**Figure 1.**
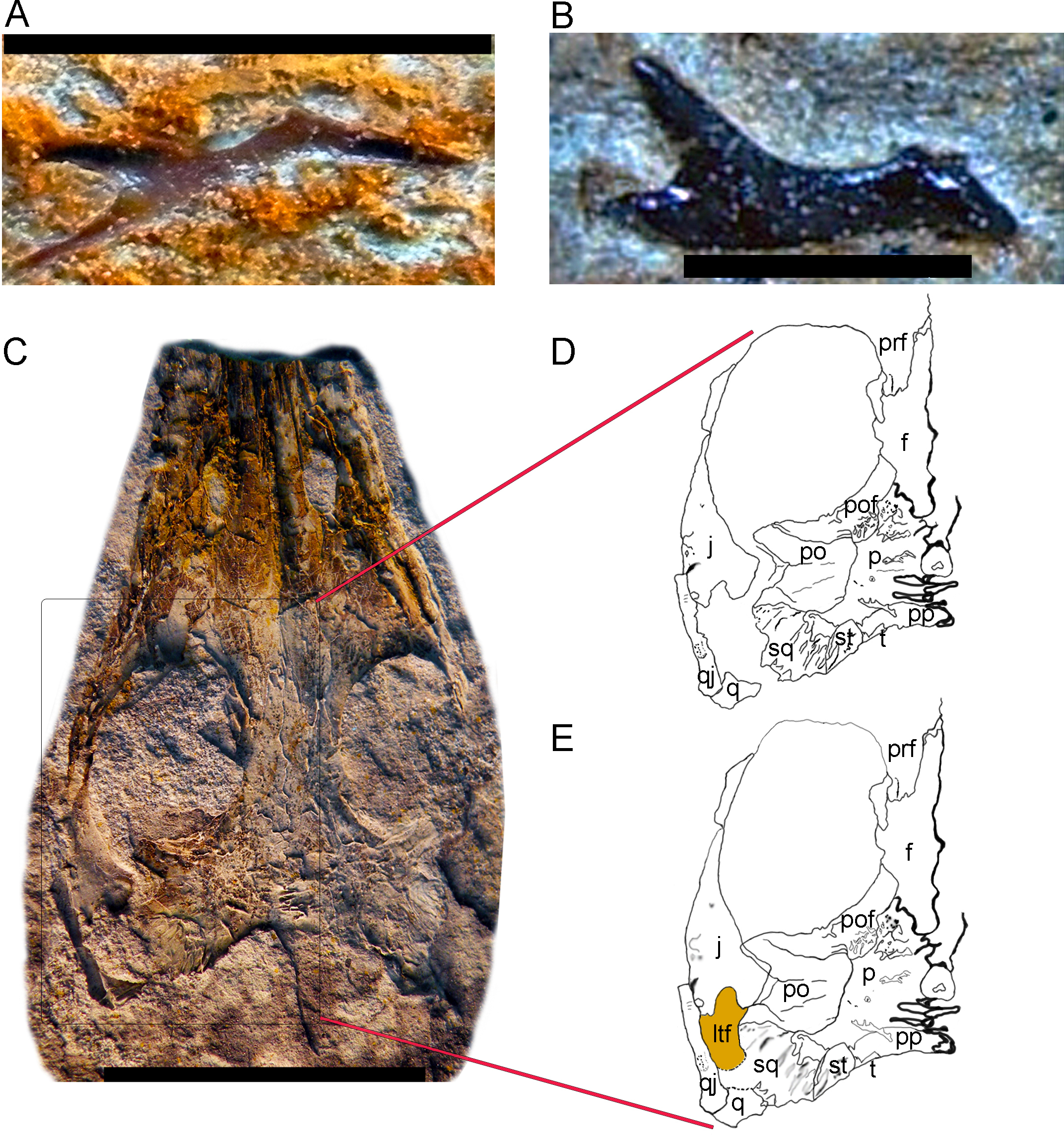
Mesosaur lateral temporal fenestra. A. FC-DPV 1462, right jugal, probably of a newborn individual showing the typical three-radiate structure normally associated with a temporal fenestra. B. FC-DPV 1083, right jugal of a juvenile individual showing the same typical three-radiate structure. C. FC-DPV 2534B, partial skull of an adult *Mesosaurus tenuidens* showing a well-preserved temporal region. Some of the bones that delimit the lateral temporal fenestra (jugal and quadratojugal) are partly disarticulated, not showing the correct anatomical configuration. However, the jugal displays the typical three-radiate structure commonly associated with the temporal fenestra. D. Schematic drawing of the FC-DPV 2534B left post orbital region as it was preserved, showing the jugal and quadratojugal partially disarticulated. E. Schematic reconstruction of the FC-DPV 2534B left post orbital region as it may have been in its natural anatomical configuration. The jugal-postorbital contact and the position of the quadratojugal relative to the jugal are approximate given that the skull bones were compressed by posterior sediment deposition, lacking their original three dimensional arrangement. However, taking into account the morphology observed in other specimens, and the general anatomy of the involved bones, the present configuration is the most plausible. Scale: A, B, 5 mm; C, 10 mm.

Aside from issues of preservation of the specimens from Uruguay compared with those of Brazil and South Africa, it is clear that some specimens from Brazil display a well-preserved temporal region featuring a lower temporal fenestra (Laurin and Piñeiro, 2017: fig. 1). Moreover, we are not the first to interpret the temporal region of *Mesosaurus tenuiden*s specimens from the Irati formation (Brazil) as displaying a lower temporal fenestra; our great predecessor von Huene (1941) illustrated, described, reconstructed, and discussed the systematic significance of that fenestra.

MacDougall et al. (2018) claim that “Laurin and Piñeiro made no effort to reexamine the *Mesosaurus* specimens that had been previously described by Modesto (2006, 2010).” This is again misleading at best; one of us (GP) examined many specimens from the collection studied by Modesto (though not the specimens that he illustrated), which were present as casts in Frankfurt and has studied good pictures of specimens from the American Museum of Natural History. Modesto has not studied the Brazilian collections containing thousand mesosaur specimens; he apparently examined materials housed in South African museums and institutions, and the best specimens from the Iratí Formation in European and North American collections. Thus, one of us (GP) has examined far more specimens of mesosaurs (79 are mentioned in our papers; see Supplementary Data Sheet 1) than Modesto (we counted only 36 specimens mentioned in his thesis and his various papers); see Supplementary Data Sheet 1 for the source of these numbers. More importantly, why has Modesto (2006) studied only one (GPIT [Institüt und Museum für Geologie und Paläontologie der Universität Tübingen] 1757-1) of the many specimens (33 are illustrated) studied by von Huene (1941) and that had led this great paleontologist and anatomist to conclude that mesosaurs had a lower temporal fenestra? The specimen (GPIT 1757-1) studied by Modesto (2006) is exposed in dorsal view and is not very informative about the temporal region. Thus, we believe that on this front, our study rests on better grounds than that of MacDougall et al. (2018).

Nevertheless, we take this opportunity to assess the impact of temporal fenestration on our analysis by analyzing our matrix again without the two temporal fenestration characters that we had (our characters 30 and 31). The search, carried out in PAUP 4.0a (build 163) for Macintosh (the latest version available as of July 28, 2018) using the branch and bound algorithm, yielded two most parsimonious tree of 328 steps (consistency index of 0.497; homoplasy index of 0.503; retention index of 0.6626) whose strict consensus is identical with the tree that we published (Laurin and Piñeiro, 2018: fig. 5). Thus, as for MacDougall et al.'s (2018) matrix, exclusion of the temporal fenestration characters does not change the results in any significant way. This refutes the claim by MacDougall et al. (2018: 5) that the variability of the temporal fenestration explains the topology that we obtained.

We also reanalyzed the matrix by MacDougall et al. (2018), using the version posted on the journal's web site, initially without any modifications. We used the heuristic search with 50 random addition sequence replicates, holding 3 trees at each step, and using the tree-bisection-reconnection (TBR) algorithm. Surprisingly, we found not 9 optimal trees of 669 steps as reported by MacDougall et al. (2018), but 225, of the same length (669 steps). The strict consensus (Fig. 2A; we illustrate it here to facilitate comparisons with our other results presented below) is identical to theirs (MacDougall et al., 2018: fig. 1A), so this might possibly attributed to how PAUP collapses or not 0-length branches or other subtle settings, but we note that their tree was rooted improperly as it implied an “anamniote” clade including *Seymouria* and diadectomorphs that excluded amniotes, whereas there is a fairly widespread consensus that diadectomorphs are more closely related to amniotes than to seymouriamorphs (e.g., Laurin and Reisz, 1995; Ruta and Coates, 2007; Marjanovic and Laurin, 2018). The rooting option of the tree is a hypothesis, not a result.

**Figure 2.**
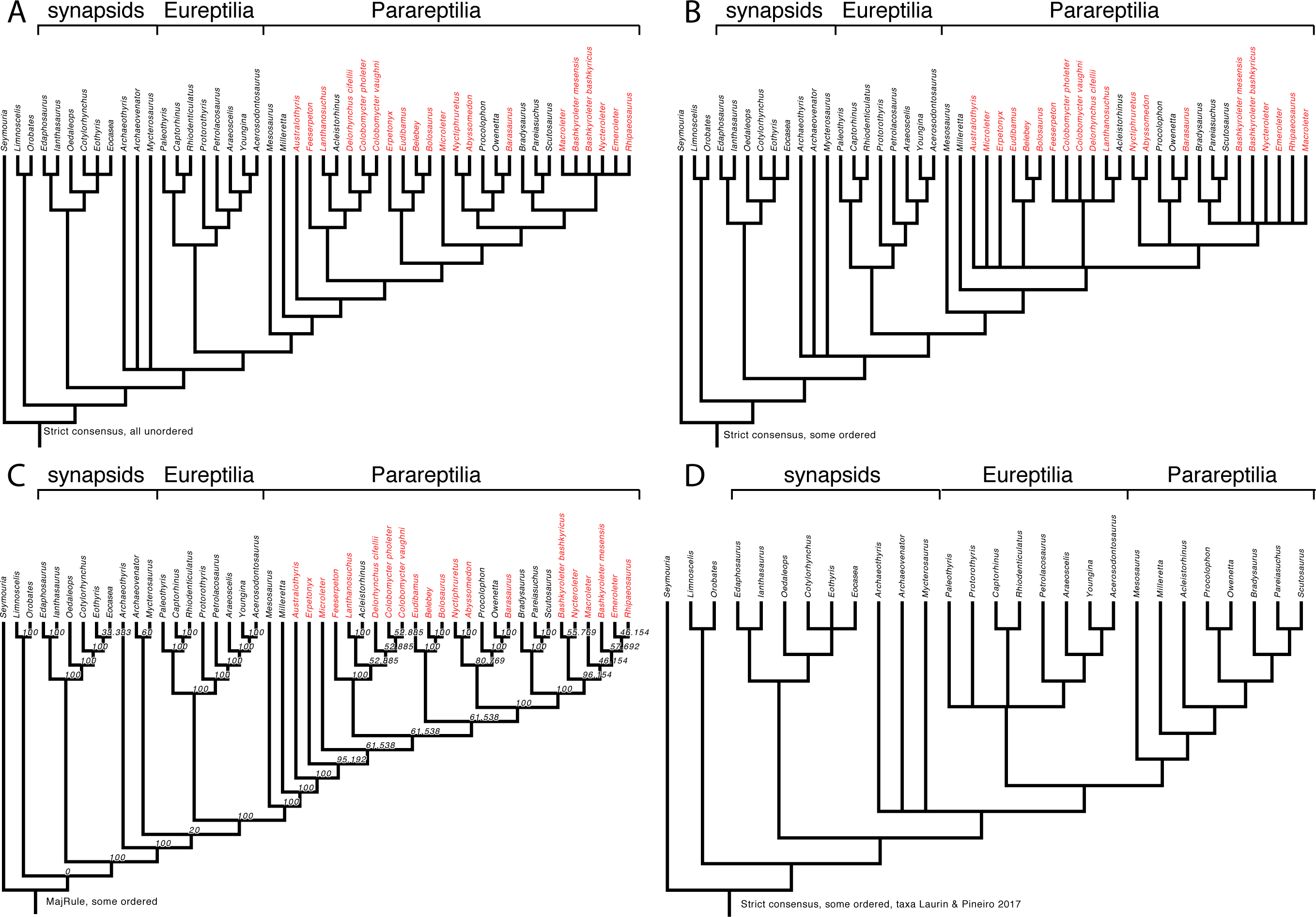
Early amniote phylogeny as assessed through various reanalyses of a slightly modified version of the matrix of MacDougall et al. (2018). (A) Strict consensus obtained by reanalyzing the data matrix without any modifications. This is basically identical with the tree obtained by MacDougall et al. (2018: fig. 1A) but it is reproduced here to facilitate comparison between the results obtained through our various analyses. (B) Strict consensus obtained by reanalyzing the data matrix after ordering characters 6, 47, 48, 52, 69, 89, 105, 129, 132, 147, 150, 165, 167, 168, 178. Note that in some cases, the order of states had to be altered to conform with the morphocline. (C) Majority-rule consensus (nodal values represent frequencies in the source trees) of the same. (D) Strict consensus obtained on the same, except that the taxa not represented in Laurin and Piñeiro (2017) were removed before analyzing the data. This refutes the claim that the topology obtained by Laurin and Piñeiro (2017) results from exclusion of several parareptile taxa (in red on parts A–C) that were considered by MacDougall et al. (2018) because exclusion of these taxa, on the matrix of MacDougall et al. (2018), does not change the topology.

MacDougall et al. (2018) did not order any characters. However, simulations have shown that for characters that form morphoclines (such as all characters that represent discretization of an inherently continuous variable, such as size or ratios between measurements), ordering states leads to better results, both in terms of power to recover true clades, and in avoiding inferring erroneous clades (Rineau et al., 2015, 2018). Thus, we have ordered the following multi-state characters (their numbering): 6, 47, 48, 52, 69, 89, 105, 129, 132, 147, 150, 165, 167, 168, 178. In some cases, we had to reorder states because the order in which they were listed made no sense if the character were to be ordered linearly, which is simplest, and which reflects the underlying quantitative (continuous) character. For instance, character 6, “Pineal foramen position: in the middle of the body of the parietal (0); displaced posteriorly (1); displaced anteriorly (2)” was reordered by inverting states 0 and 1. Characters 47, 89, 147, and 178 were likewise reordered. Character 147, about humeral morphology, requires an explanation. MacDougall et al. (2018: supplementary data sheet 1) indicate that this is “Modified from Laurin and Reisz, 1995 #104”. The only modification that we see is the suppression of the ratio that served to make state definitions more objective. In Laurin and Reisz (1995), we had indicated that the states depended on the ratio between distal humeral head width and humeral length, using 35% and 65% as thresholds. With this information, it is clear that the states should be ordered. In the list of characters (see Supplementary Data Sheet 2), we have tracked the changes made to state numbering to facilitate comparisons between our settings and those used by MacDougall et al. (2018). We also provide the revised data matrix (Supplementary Data Sheet 3) with the states reordered and ordering (or not) of multi-state characters specified, to facilitate its use by other scientists.

The search on the matrix with some characters ordered (as mentioned above), conducted with the same settings as mentioned above, yielded 1560 trees requiring 679 steps each and with a consistency index of 0.2975, a homoplasy index of 0.7025, and a retention index of 0.6446. Their strict consensus (Fig. 2B), unsurprisingly, is less resolved than the tree reported by MacDougall et al. (2018: fig. 1A). Notably, there is a large polytomy near the base of Parareptilia, in the clade that includes all parareptiles except for *Milleretta,* another one at the base of Lanthanosuchoidea, and yet another polytomy is formed by Nyctiphruretidae, Procolophonoidea, and a clade that includes Pareiasauria and nycteroleterids. In the latter, Nycteroleteridae is possibly paraphyletic as its component taxa form a polytomy that also includes Pareiasauria. This last polytomy was also obtained by MacDougall et al. (2018: fig. 1B) when they excluded the characters linked with temporal fenestration, so the monophyly of Nycteroleteridae might be worth reassessing. It might be tempting to view the lower resolution of our tree as a refutation of our ordering scheme, but it is more likely that the clades found by MacDougall et al. (2018: fig. 1A) are erroneous because simulations have shown that not ordering intrinsically ordered characters yields a greater proportion of erroneous clades (Rineau et al., 2015, 2018).

The majority-rule consensus tree of the 1560 trees obtained with some states ordered (Fig. 2C) reveals some interesting information not provided by MacDougall et al. (2018). Namely, varanopids may be more closely related to other sauropsids than to *Archaeothyris*. Support for this is admittedly very low (20% of the most parsimonious trees), but it is also found (with the same frequency) when no states are ordered (not shown here, but available in Supplementary Data 3).

MacDougall et al. (2018: 5) claimed that the omission of several parareptile taxa in our matrix explained our topology (with mesosaurs outside Parareptilia). To test this hypothesis, we deleted from their matrix the parareptiles that they included but that were excluded from our matrix. In this test, we did not remove low-ranking taxa that are part of higher-ranking taxa that we included. Thus, the OTUs belonging to Pareiasauria included in MacDougall et al. (2018) were retained, but we excluded *Australothyris, Microleter, Nyctiphruretus, Barasaurus, Bashkyroleter mesensis, Bashkyroleter bashkyricus, Nycteroleter, Emeroleter, Ripaeosaurus, Lanthanosuchus, Feeserpeton, Colobomycter pholeter, Colobomycter vaughni, Delorhynchus cifellii, Abyssomedon, Eudibamus, Belebey, Erpetonyx, and Bolosaurus*. The search, conducted with the same settings as above, yielded 18 trees of 465 steps, and with a consistency index of 0.4151, a homoplasy index of 0.5849, and a retention index of 0.6397. Their strict consensus resembles closely the tree published by MacDougall et al. (2018: fig. 1A), with *Mesosaurus* as the basalmost parareptile, and varanopids and the presumed ophiacodontid *Archaeothyris* appearing at the base of Sauropsida (Fig. 2D). We also repeated this analysis with the characters linked to temporal fenestration excluded, as in MacDougall et al.'s (2018) second analysis. The results (not shown, but available in Supplementary Data Sheet 4) show a very similar topology, with mesosaurs at the base of Parareptilia, varanopids and *Archaeothyris* forming a polytomy at the base of Sauropsida, etc. This, along with the fact that exclusion of the temporal fenestration characters from our own matrix does not change the resulting topology, directly refutes the conclusion by MacDougall et al. (2018: 5) that “we illustrate that the lack of taxa in their matrix combined with the variability of temporal fenestration in Reptilia are likely contributing to the tree topology that they obtained in their phylogenetic analysis…”. Obviously, the differences in topologies (including about the position of mesosaurs) supported by our matrix and theirs must rather be attributed to changes in character scores and character selection.

## 5 Discussion

The differences in character treatment mentioned above (including the decision to order or not the states, the order of the states, and more importantly, the way in which states are delimited) explains partly why we did not wish to use, as the basis on which to build our matrix, more recent versions of the same matrix, which were produced in Reisz' lab after one of us (ML) left that lab, or other matrices with much more tenuous links with that matrix (e.g., Schoch and Sues, 2017 [Schoch, 2017 #22813]). In addition to the minor problems discussed above, our experience has convinced us that it is very difficult for a systematist to expand another systematist's matrix because a certain amount of subjective judgment is involved in character scoring, and each systematist has his own perception of a given character state. The only way to ensure coherent application of character step definitions is for a single systematist to score a given character for all taxa included in a matrix. Thus, to adequately reuse another systematis' matrix, it is unfortunately necessary to more or less redo all the scoring, after having checked, as we did, which character should be ordered and how. And of course, is it desirable to have studied several specimens directly, as illustrations only provide part of the information, even though recent 3D imaging developments led to improvements on that front (e.g. Tissier et al., 2017 [Tissier, 2017 #22863]).

MacDougall et al. (2018) argue that recent works have attempted to improve on state definitions to solve these problems but our own examination of recently-published data matrices leads us to believe that we are unfortunately still very far from that ideal situation. In fact, some of the changes introduced by MacDougall et al. (2018), as for their character 147 (humeral morphology), by removing the ratio and values used as thresholds by Laurin and Reisz (1995), make the character scoring less replicable, so their work certainly does not back up their claim that substantial progress has been made on this front (at least, in the last two decades). Updating other systematists matrices is extremely time-consuming, if it is done carefully, as illustrated by the reanalysis of the matrix by Ruta and Coates (2007), initiated by one of us (ML) and first tackled in the doctoral thesis of Germain (2008), continued in another doctoral thesis (Marjanović, 2010), and recently published as a pre-print, in two successive versions (Marjanović and Laurin, 2015, 2018). This work has been under review since 2015 and will be published soon. Even though this is an extreme case with respect to the time investment it required, it does not represent the most thorough rescoring of a matrix that we have performed. Rather, our work on a much smaller data matrix of 23 taxa and 42 characters, in which it was feasible to re-examine the scoring of all cells, resulted in changes in the score of 35% of the cells of the matrix, spread over all taxa and all but two characters (Laurin and Marjanovic, 2008). These examples illustrate well the point that reusing other people's data matrices is not a trivial exercise, at least if it is to be done correctly, contrary to MacDougall et al.'s (2018) claim. By starting from one of our own matrices, we minimized this problem, without eliminating it completely. According to MaShea (2000: 330), “It has been said that most scientists would rather use another scientist's toothbrush than his terminology.” We feel the same about other author's phenotypic data matrices (and the stakes are much higher), unless time is taken to study thoroughly each character and its distribution to ensure that whoever reworks a matrix has the same understanding about each character state. To sum up, our method did not “patently ignore over two decades of parareptilian research”, and the resulting matrix is not outdated; it results from a deliberate choice to obtain a reasonably reliable matrix with an appropriate taxonomic sample for a reasonable time investment. It is a compromise, and as such, we recognize that it is imperfect and that it can be improved; we only disagree about how this should be done. Our preferred path is to work with a matrix that we know well and expand it ourselves, rather than rely on a patchwork containing additions by several scientists who may not have understood his predecessor's concept of each character.

We showed, through the new analyses presented above, that taxon selection and temporal fenestration variability do not appear to explain the differences in topologies obtained by our group and MacDougall et al. (2018). Strangely, MacDougall et al. (2018) had all the data required to check these claims themselves but did not bother to perform the necessary analyses before publishing their conclusions. At least one other reasonably recent phylogenetic analysis recovered mesosaurs in a fairly basal (though unresolved) position among amniotes (Hill, 2005), outside Parareptilia.

MacDougall et al. (2018) also failed to recognize that differences between our taxon selection and theirs reflect different strategies and goals. Rather than trying to sample densely parareptiles, we included three turtle OTUs (*Odontochelys, Proganochelys* and Chelonii) because they may be parareptiles (Laurin and Reisz, 1995) and because the presence of this extant taxon in the matrix could have altered the most parsimonious topology, given their strongly divergent morphology (compared to parareptiles and diapsids). Thus, to minimize the risks of getting phylogenetic artifacts, including turtles may be more important than adding more parareptile taxa that resemble more the other parareptile taxa already present in our matrix, though the taxon deletion tests that we carried out (Laurin and Piñeiro, 2017) neither confirm nor refute this. Turtle origins remains one of the great controversies of vertebrate phylogeny (e.g. Lyson et al., 2010, 2013; Lee, 2013), which is potentially relevant to several zoologists, and our matrix, which we will continue to develop by adding taxa and characters (through the project involving I. Werneburg, G. Ferreira and M. Rabi evoked above), is a step in that direction. MacDougall et al. (2018) built their matrix for a more limited goal (not looking beyond Permo-Triassic taxa); both approaches are valid and complementary. Of course, it is possible to include both many additional parareptile taxa and turtles, but the more taxa are added, the more characters need to be included to resolve their relationships, and given that research time is limited, the more cells a matrix includes, the least time goes into scoring each of them. Thus, there is an optimal number of taxa and characters for phenotypic matrices; more is not always better.

The fact that six authors collaborated to publish the short paper (MacDougall et al., 2018) to respond to our paper (which has only two co-authors) might be interpreted as strengthening that paper by showing that a significant proportion of the (very small) community of experts on Permo-Carboniferous amniotes agreed sufficiently with each other to write that paper. However, majority opinion has never been a safe guide for scientific progress, as illustrated by the pamphlet “100 Authors against Einstein”, which attempted to refute Einstein's theory of relativity. Einstein reportedly replied (Hawking, 1993: 98) “If I were wrong, then one [author] would have been enough!”

## 6 Acknowledgments

We thank Mark MacDougall for sending the draft of the paper and the supplements before their publication (but after definitive acceptance of their paper), at our request.

## 7 Author Contributions Statement

ML planned this research, carried it out and wrote most of the draft. GP wrote part of the text, drafted figure 2, and provided comments to improve other parts of the text.

## 11 Supplementary Material

The Supplementary Material for this article can be found online on the bioRxiv web site.

